# Resolving interface structure and local internal mechanics of mitotic chromosomes

**DOI:** 10.1101/2024.08.20.608279

**Authors:** Andrea Ridolfi, Hannes Witt, Janni Harju, Tinka V. M. Clement, Erwin E. J. G. Peterman, Chase P. Broedersz, Gijs J. L. Wuite

## Abstract

The interface of chromosomes enables them to interact with the cell environment and is crucial for their mechanical stability during mitosis. Here, we use Atomic Force Microscopy (AFM) to probe the interface and local micromechanics of the highly condensed and complex chromatin network of native human mitotic chromosomes. Our AFM images provide detailed snapshots of chromatin loops and Sister-Chromatids Intertwines. A scaling analysis of these images reveals that the chromatin surface has fractal nature. AFM-based Force Spectroscopy and microrheology further show that chromosomes can resist severe deformations, elastically recovering their initial shape following two characteristic timescales. Localized indentations over the chromatids reveal that the spatially varying micromechanics of the chromatin network is largely governed by chromatin density. Together, our AFM investigation provides new insights into the structure and local mechanics of mitotic chromosomes, offering a toolbox for further characterization of complex biological structures, such as chromosomes, down to the nanoscale.

---

During mitosis, the chromatin of eukaryotic cells undergoes a dramatic reorganization that results in the formation of highly condensed rod-like structures known as mitotic chromosomes ^1,2^. In this process, chromosomes experience exogenous forces from the action of the mitotic spindle, which first pushes chromosomes towards the metaphase plate and then pulls the sister chromatids apart towards the spindle poles^2–4^. How chromosomes interact with their external environment and withstand these local forces is still unclear, even though this represents a key step for the successful completion of the mitotic cycle. In this context, AFM offers the unique possibility of accessing both the interface structure and the local micromechanics of mitotic chromosomes, thereby allowing a thorough characterization of their chromatin architecture.

Current structural models ^5^ show that mitotic chromosomes consist of chromatin loops extruded from a central protein-rich scaffold where most of the condensin II protein complexes are located. Condensin I complexes further compact each condensin II-mediated loop into smaller loops. Numerous techniques were used to probe the whole structure of mitotic chromosomes ^6–10^; however, the high susceptibility of chromosomes to external conditions^11,12^ and the soft yet very dense nature of their chromatin architecture^8^ still complicate access to smaller (sub-micron) features of the chromosome architecture and to their mechanics. Micropipette-based manipulation or optical tweezers (OTs) can be used to probe the macroscopic mechanical response of chromosomes to stretching deformations ^11,13–15^ but are unable to access the local mechanics of the chromatin network at the nanoscale level. This knowledge gap can be bridged by exploiting the capabilities of Atomic Force Microscopy (AFM).

Indeed, AFM can be used to characterize the interface structure of mitotic chromosomes over the entire range of its length. In addition, AFM can be used to probe chromosome mechanics, via AFM-based Force Spectroscopy (AFM-FS)^16^, and microrheology, both with high spatial accuracy. All these measurements can be performed in a liquid environment, mimicking the native mitotic conditions. AFM has been applied to chromosomes before, though in most cases on fixed or dried samples. The few AFM studies characterizing chromosomes in liquid conditions have employed PBS^17,18^ or water as imaging solutions^19^, potentially altering both chromosome structure and mechanics due to the susceptibility of the chromatin network to variations in the physico-chemical properties of the environment ^11,12^. Furthermore, most of these studies used chromosome spreads ^18–21^, where chromosomes are covered by cytoplasmic debris that can only be removed via specific enzymatic treatments that could alter the delicate chromatin network ^22^.

Here, we show how AFM can be used to resolve the interface structure and the internal mechanics of native human mitotic chromosomes in a liquid solution mimicking the physico-chemical properties of cells in mitosis. Our AFM images probe the chromosome interface over the entire range of scales of its structural hierarchy, revealing features like chromatin loops and Sister Chromatids Intertwines (SCIs). From the image analysis, we find that chromatin does not exhibit a specific higher-order organization and that the chromosome surface possesses a statistical fractal nature with self-similar scaling. Results from AFM-FS show that chromosomes can withstand and elastically recover from extremely high forces. Complementing these results with microrheology experiments, we find that the chromatin network relaxes on two prominent time scales. Finally, by performing localized indentations over the entire width of the chromatids we reveal that the local stiffness of the chromatin network is highly variable, both along a single chromosome and among different chromosomes. The variable micromechanics is mostly mediated by local chromatin density.

## AFM imaging of native mitotic chromosomes under physiological solution

The properties of the liquid environment in which mitotic chromosomes are probed play a crucial role on their structure and mechanics ^11,12^. Here, isolated mitotic chromosomes were probed within a polyamine (PA) buffer that mimics the physico-chemical conditions of the mitotic environment in cells ^23^ (Methods). In these conditions, mitotic chromosomes are fully hydrated and appear as a soft network of highly entangled chromatin fibers, displaying structural hierarchy at different length-scales (Figure 1a, b and c). With our chromosomes, the sister chromatids are in close contact and attached to each other at several points along their length by chromatin fibers (Figure 1b). Blue arrows in Figure 1b point at several chromatin loops stemming from inside the chromosome body; these features support the loop arrangement proposed by Gibcus et al. ^5^, even though the overall chromatin network seems more disordered and randomly coiled than the predicted bottlebrush with a layered organization of loops. This is also supported by the presence of fibers not arranged in loops and spanning distances greater than the typical loop size (white arrows in Figure 1b). Figure 1c highlights two different types of SCIs: those formed by individual chromatin fibers connected to both sister chromatids and those consisting of interlaced fibers forming loop structures (Figure 1c and its schematic illustration in Figure 1d). The presence of SCIs is in line with earlier studies demonstrating that also nocodazole-arrested yeast cells present these topological links ^24^. Despite the high resolution of Figure 1b and c, the displayed features appear not completely ‘sharp’; we interpret this observation as an indication that our imaging conditions still allow for some thermal motion of the soft chromatin architecture.

**Figure 1:**
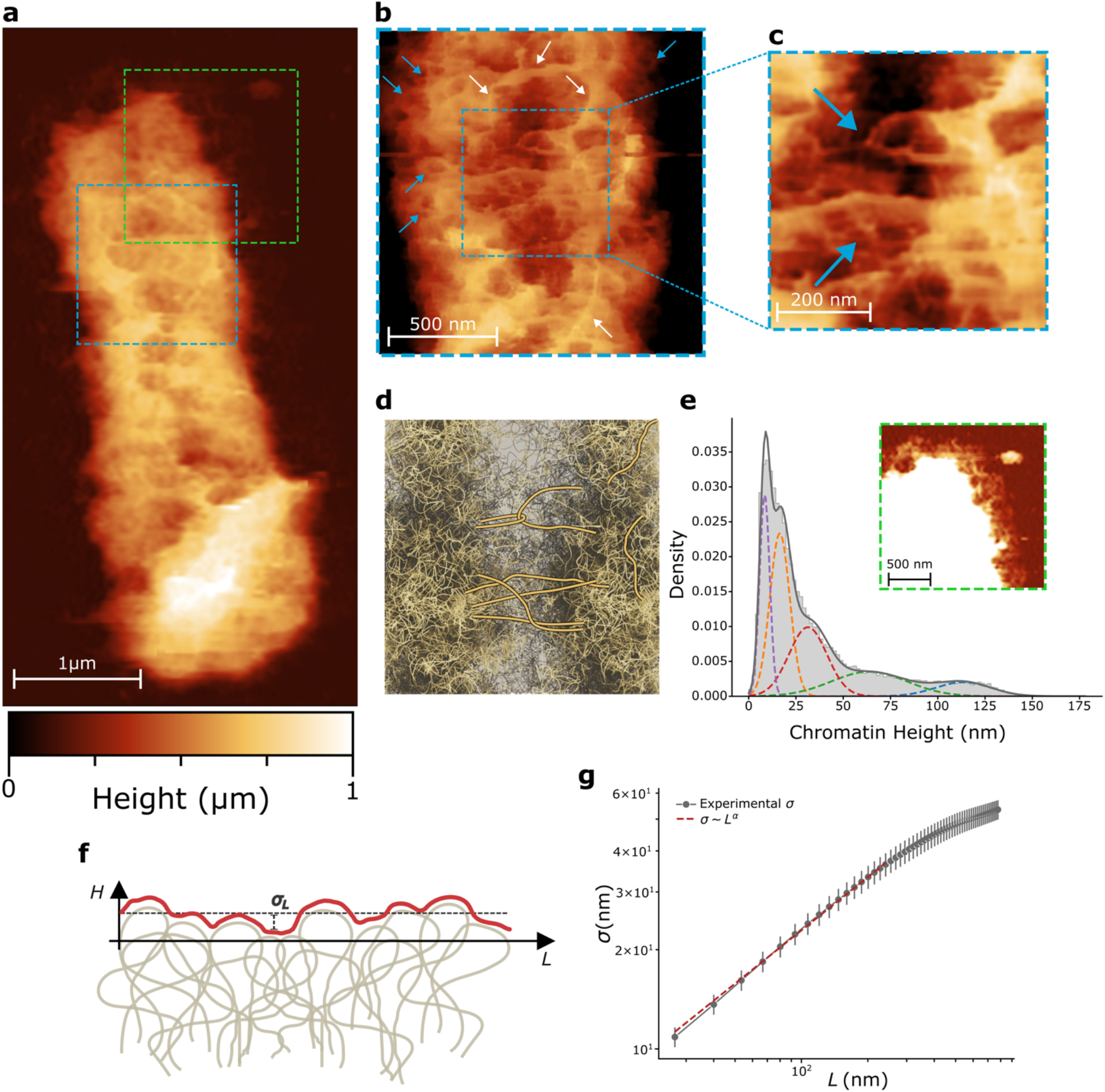
AFM imaging of native human mitotic chromosomes. a) AFM topography of a single native human mitotic chromosome imaged within PA buffer. b) AFM topographical image zooming in the area within the blue dashed square reported in a); blue arrows point to loop structures stemming from the central part of each chromatid while white arrows indicate chromatin fibers not arranged in loop-like structures. c) Inset from b) displaying different types of SCIs holding together the two sister chromatids; the upper arrow points at two loops interlaced together while the lower one shows that multiple chromatin fibers are still connected with both sister chromatids. d) Schematic representation of c), highlighting the two types of SCIs and two randomly coiled fibers. e) The inset shows a high-contrast image of the area within the green dashed square in a), in these conditions it is possible to see the halo of chromatin surrounding the chromosome (full picture in Figure S1). The histogram describes the height distribution of the chromatin halo, fitted with a combination of 5 Gaussian curves (the minimum number that yielded an accurate fit). f) Illustrative height profile (red trace) describing the height fluctuations of the chromosome surface (gray structure). The horizontal dashed black line indicates the mean surface height. The standard deviation (σ_L_) characterizes the surface roughness. The magnitude of σ_L_ depends on the sampled length scale L. g) Averaging all the values of σ_L_ along different longitudinal profiles yields a measure of the average surface roughness σ_L_; repeating the procedure for different length scales L reveals a power-law scaling regime (dashed red line), indicating self-affinity.

### Chromatin organization and the fractal nature of chromosome surface

In the higher contrast, zoomed in image in the inset of Figure 1e and S1, chromatin fibers and loops protruding outwards and forming a “halo” around the chromosome body can be observed. Similar structures have also been reported before ^19^ and have been attributed to partial unraveling of the chromosome during the sample preparation and adsorption steps. In our images the “chromatin halo” appears approximately four times smaller, which might reflect the gentler sample preparation. In Figure 1e the height distribution of the chromatin halo (Figure S1) is shown. The distribution is fitted with a sum of 5 Gaussian functions, where the first peak is located at approximately 8.5 nm, in agreement with previous studies^6,25^ and compatible with the “beads on a string” model for chromatin organization. The centers of the other Gaussian functions are located at 16.5, 31.3, 63.7, and 113.4 nm, all approximately multiples of the main 8.5 nm peak, probably describing the stacking of multiple chromatin fibers. The absence of prominent secondary peaks in the height distribution suggests that there is no preferential higher order organization for chromatin in the outer layers of the chromosome.

To gain more insights into the nature of the chromosome surface, we analyzed its roughness. A natural way to characterize the roughness of a random surface is to explore its fractal nature. This can be done by analyzing the roughness at different length-scales to test for self-affinity ^26,27^. Briefly, we traced multiple height profiles both parallel and perpendicular to the long axis of the chromosome (Figure S2). Along each trace, we then measured the standard deviation of the height, *σ*, which characterizes the surface roughness of the chromosome (Figure 1f). A regular non-fractal interface would exhibit a roughness that is independent of the length scale over which it is determined, provided this length scale exceeds the largest associated correlation length in the system. By contrast, for mitotic chromosomes plotting the standard deviation as a function of length-scale *L* (Figure 1g, Figure S3) reveals a power-law scaling regime *σ* ∼ *L*^*α*^ for length-scales from 40 to 250 nm, with *α* = 0.6 ± 0.1 (value from fit ± standard error) for longitudinal (Figure 1g) and a similar *α* = 0.6 ± 0.1 (value from fit ± standard error) for perpendicular traces (Figure S3). Such scaling relations indicate a non-trivial self-similarity that controls the roughness over a broad range of length scales.

### Chromosomes elastically recover from repeated high-force deformations

Being able to access the native structure of mitotic chromosomes allows leveraging AFM-based Force Spectroscopy (AFM-FS) to probe local chromosome mechanics. AFM-FS employs higher forces than AFM imaging, potentially altering the chromatin architecture and its stability. To rule out such potential artifacts, we performed repeated indentations at the same location on the chromosome and monitored the variation of different structural and mechanical descriptors. Each AFM-FS measurement results in a force curve representing the force experienced by the tip as it penetrates and retracts from the sample^16^. In the representative force curve of Figure 2a, the red cross indicates the contact point at which the AFM tip starts to push against the chromosome and can be interpreted as an indirect measure of the sample height. Figure 2b shows that after 30 repeated indentations (for 4 different chromosomes), the contact point (i.e., chromosome height) varies less than 5% from its initial value. This result rules out the introduction of any artifact from the probing process and highlights that chromosomes can withstand and elastically recover from local indentations extending deep through their entire body and reaching forces up to about 10 nN.

**Figure 2:**
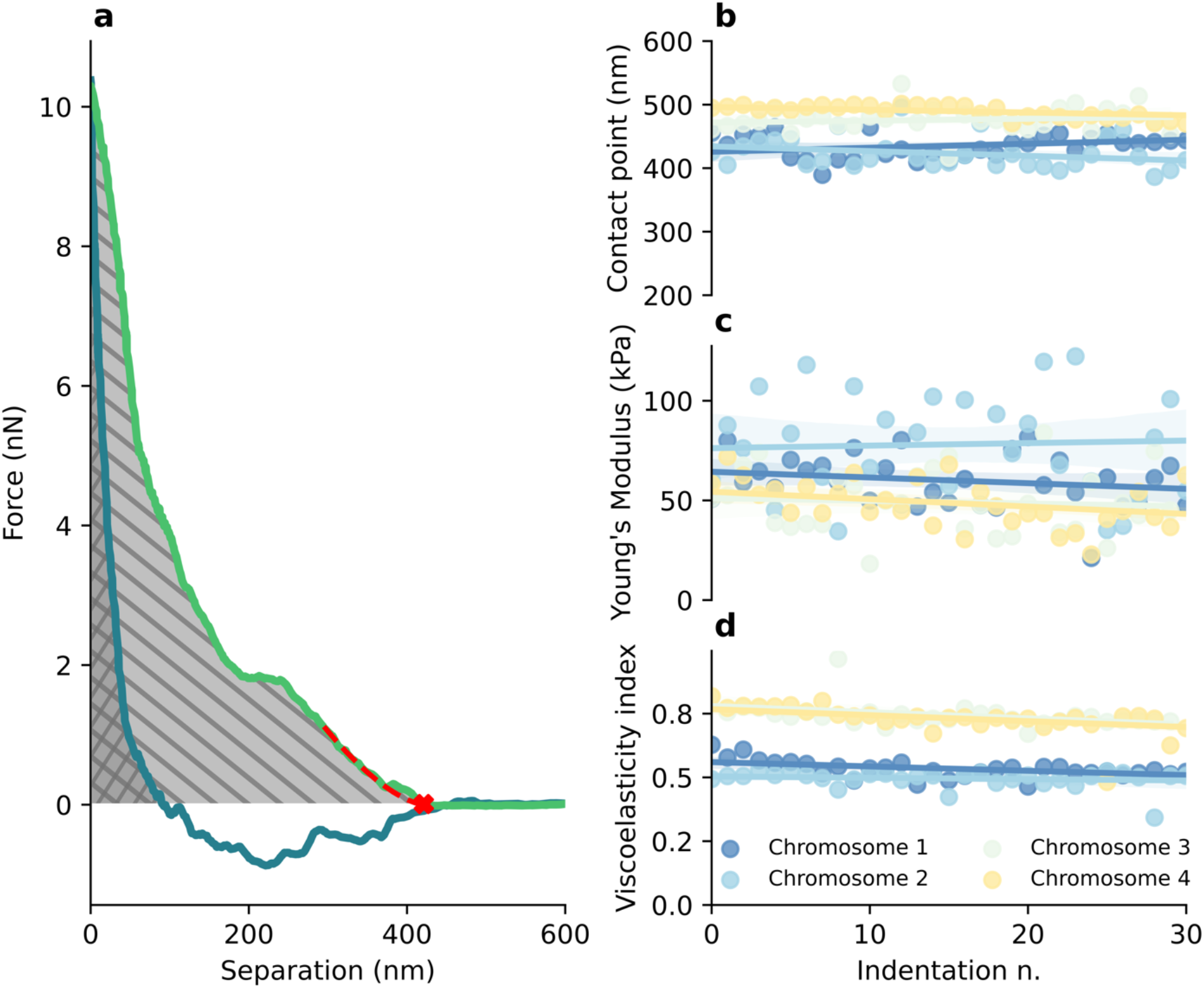
a) Representative force curve from an AFM-FS measurement; the green trace describes the indenting curve while the dark blue trace the retracting one. The red cross identifies the contact point, where the AFM tip starts to push against the chromosome body while the red dashed line is the fit used to calculate the Young’s Modulus. Back-diagonal and diagonal hatch patterns identify the areas A_1_and A_2_, under the indenting and retracting curves during the loading and unloading regimes, respectively (i.e., where the tip-sample interactions result in positive force values). These areas are used to calculate the viscoelasticity index, defined as η = 1 – A_2_⁄A_1_. b) Contact point, c) Young’s Modulus and d) Viscoelasticity index values obtained by performing 30 repeated indentations on 4 different chromosomes. Values for each point were estimated from the single force curves; shaded areas represent the 95% confidence intervals.

From the same force curves, it is possible to extract mechanical descriptors of the elastic and inelastic response of chromosomes to the applied deformations. The Young’s Modulus, which quantifies the relationship between the stress and the strain applied during the indentation process within the linear elastic regime, was obtained by fitting the initial part of the indenting force curve with a modified Hertz model ^28^ (red dashed line in Figure 2a). The inelastic mechanical response, on the other hand, is related to the energy dissipated during the indentation, due to viscous deformations. This can be described by the so-called *viscoelasticity index, η*^29,30^ (caption of Figure 2a). A purely elastic response would result in *η* = 0, while *η* = 1 indicates a purely viscous response. The traces reported in Figure 2c and d show that both the Young’s Modulus and viscoelasticity index displayed negligible variation during the four repeated-indentation series, indicating that chromosome mechanics is not affected by deformation history. Furthermore, the individual viscoelasticity index values (Figure 2d), ranging from 0.5 to 0.8, indicate that, between the loading and unloading regime, a fraction of the indentation energy is dissipated.

### Chromatin is responsible for the fluid-like behavior of chromosomes

Further insights into the energy dissipation and time relaxation of the chromatin network can be obtained by leveraging AFM-based microrheology ^31^.To this aim, we used a spherical probe with a radius of 870 nm (covering large portions of the chromosome body) to apply oscillatory deformations to the chromosomes. Figure 3a shows a representative microrheology force-time trace, where the tip is brought in contact with the chromosome and then oscillated at specific frequencies (Methods). Probing chromosome response to oscillatory deformations allowed for the calculation of the Shear Storage Modulus (G’) and the Shear Loss Modulus (G’’), which give insights in the elastic and viscous responses of chromosomes at different deformation rates. Figure 3b and c report the values of G’ and G’’ obtained at different oscillation frequencies for 19 chromosomes. Although the values vary by almost an order of magnitude between different chromosomes, both G’ and G’’ show a highly consistent frequency dependence. Averaged data in Figure 3d show that both G’ and G’’ increase with increasing frequency. G’’ has a stronger frequency dependence than G’, even though, within the probed frequency range, the elastic component (G’) remains dominant. This is also reflected by the frequency-dependent loss tangent (Figure 3e), which is defined as the ratio between G’’ and G’ and quantifies the dissipative contributions in the sample (a loss tangent of zero indicates a completely elastic response). Furthermore, both G’ and G’’ exhibit a frequency dependence consistent with *ω*^1/2^ (dashed gray trace) at large frequencies, corresponding to the high-frequency regime of the Rouse model for solutions of flexible chains^32^.

**Figure 3:**
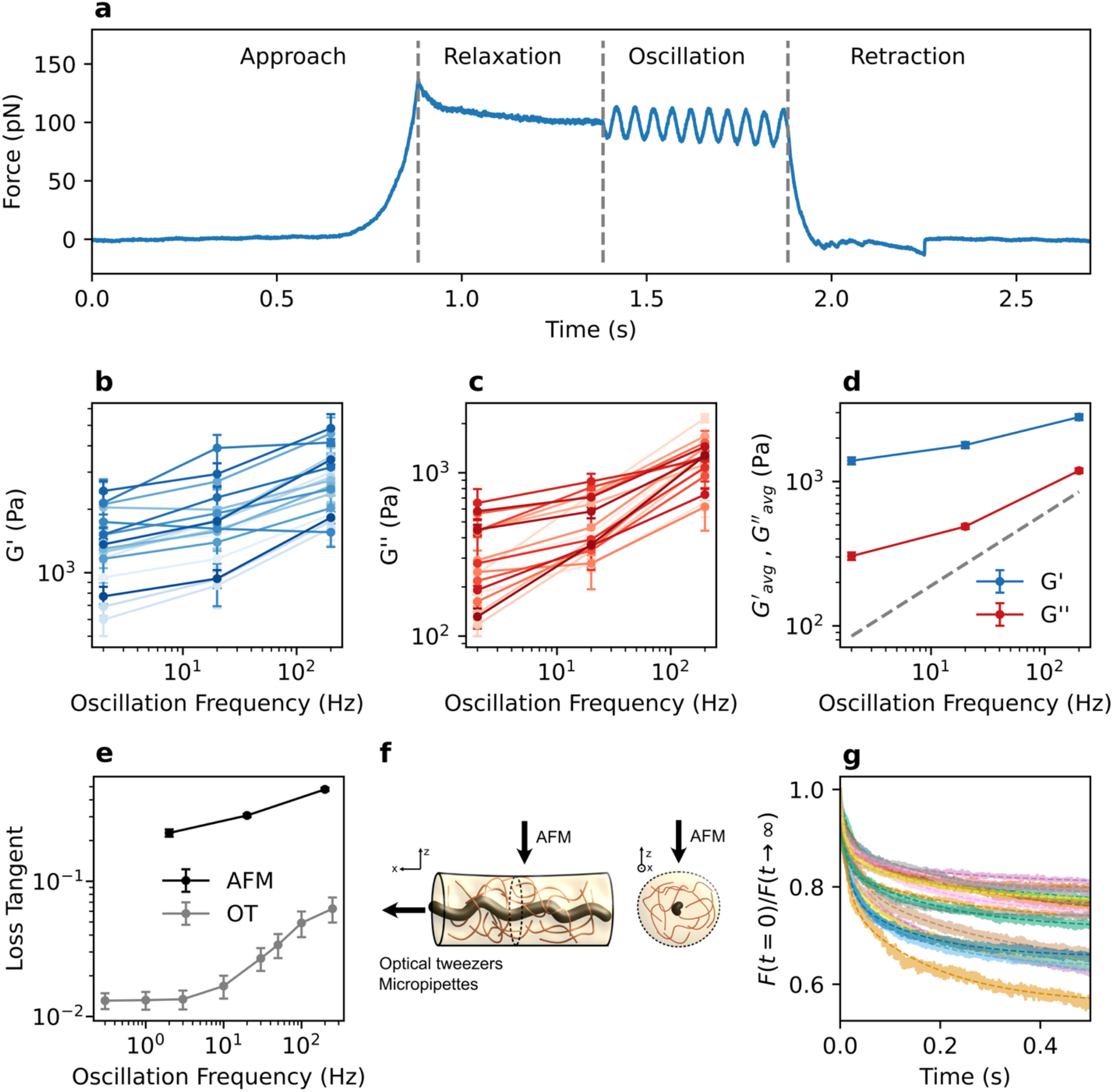
AFM-based microrheology on native human mitotic chromosomes. a) Representative force curve (plotted as a function of time) obtained from a single measurement; the tip approaches the sample and penetrate until a force of approximately 150 pN is reached. After that, the tip is kept at a constant height for 0.5 s while the chromosome structure relaxes from the initial deformation. Subsequently, the tip is oscillated for 10 periods of time and then retracted from the chromosome. Measurements were repeated multiple times on different locations in order to probe the entire chromosome body. For each chromosome this procedure was repeated using three different oscillatory frequencies, 2, 20 and 200 Hz. The oscillatory measurements allowed calculating the Shear Storage Modulus (G’) and the Shear Loss Modulus, for the three oscillatory frequencies, b) and c) respectively. Each point in b) and c) represents the average G’ and G’’ obtained from all the measurements performed on a specific chromosome; error bars are the respective SEM. d) Values of G’ and G’’ averaged over all the probed chromosomes and plotted as a function of the oscillation frequency; error bars are the SEM. The dashed gray line indicates a scaling of ω^1/2^, corresponding to the high-frequency regime of the Rouse model. e) Loss tangent as a function of the oscillation frequency; the black trace is obtained from the values of G’ and G’’ shown in d) while the gray trace reports the values obtained by Meijering et al.^15^ using OT. f) Schematics showing that AFM operates perpendicularly to the chromosome long axis, hence mainly probing the mechanical response of the chromatin network. OTs and micropipettes operate parallel to the chromosome long axis, probing the contributions of the chromatin network and the protein-rich central scaffold in parallel. g) Average FRCs for the probed chromosomes, each curve is obtained by averaging the “Relaxation” part of all the curves performed on a single chromosome. The dashed lines describe the double exponential function (Equation 1) used to fit the curves.

In the same figure, data obtained from the AFM (black trace) are compared with data obtained using OT^15^ (gray trace). For both methods, the loss tangent is dependent on the oscillation frequency; however, values obtained with AFM are almost one order of magnitude larger than those obtained with OT. This indicates that the chromosome behaves much more fluidly when deformed using AFM perpendicularly to its long axis than when stretched along the long axis using OT (Figure 3f). Earlier, we have shown that the viscoelastic response of chromosomes along the long axis can be modeled to arise from an elastic central scaffold in *parallel* with an effective solution of entangled flexible fibers, representing chromatin loops dangling around the central scaffold^11^ (Figure 3f). Based on this model, it is reasonable to assume that in our measurements the AFM tip is probing the same structures in *series*. Given the shallow indentation characterizing the Approach phase and the small amplitude of the oscillations (10 nm), the observed fluid-like behavior of the chromosome can then be understood as arising solely from the response of the entangled chromatin loops, with minimal contribution from the stiffer protein-rich scaffold ^2,5,33^.

### The chromatin network recovers from deformations following two characteristic timescales

Further analysis of the Force Relaxation Curves (FRCs, the Relaxation phase in Figure 3a) can provide additional information on how the chromatin network relaxes from the applied deformations. As shown by previous studies^34^, FRCs can be fitted with a double exponential function:

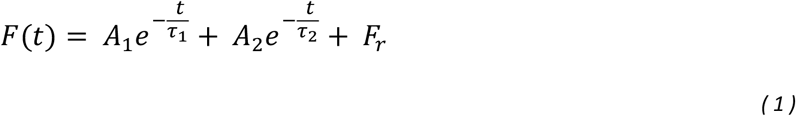

Where *A*_1_, *A*_2_ are the amplitudes and *τ*_1_, *τ*_2_, the timescales of the fast and slow relaxation processes, respectively, and *F*_*r*_ = *F*(*t* → ∞). Fitting the average FRCs of Figure 3g yielded an average *τ*_1_ of 0.014 ± 0.001 s and *τ*_2_ of 0.20 ± 0.02 s (mean ± SEM). Despite the variations in the values of the force drops (i.e., *F*(0) – *F*_*r*_) among the probed chromosomes (Table S1), neither *τ*_1_ nor *τ*_2_ showed any significant force-drop dependence (Figure S4), suggesting that the initial deformation is within the linear elastic regime. Micropipette experiments have been used before to probe the force-relaxation response of mitotic chromosomes along their long axis (Figure 3f), obtaining a characteristic timescale of 2 s^13^. The slower relaxation time during micropipette stretching could be indicative of larger structural re-arrangements involving the entire chromosome body. On the other hand, the faster relaxation behavior observed with AFM most likely accounts for only minor rearrangements of the chromatin architecture perpendicular to the chromosome long axis. In the same micropipette study ^13^ it was also found that the chromosome structure recovers the initial width (after being stretched) in approximately 0.05 s; this faster timescale is comparable with the value of *τ*_1_, which could hence describe the flow of liquid back into the chromatin network. The slower timescale, *τ*_2_, would then account for the structural reorganization of the chromatin network in the direction perpendicular to the long axis.

### Chromatin density mediates chromatid’s mechanics

According to the nested loop model ^5^ and electron-microscopy images^35^ the density of the chromatin network is not uniform, which could cause the micromechanical properties to vary within the chromatid cross-section. Figure 4a shows that the presence of both condensin I and II complexes in the central part of each chromatid^9^ results in a higher density of chromatin compared to the outer layers. The overlapping portions in between the two sister chromatids present additional chromatin entanglements and overlaps, likely produced by SCIs. To test whether these cross-chromatid structural heterogeneities translate into different mechanical responses, we performed indentations along lines perpendicular to the chromosome central axis (CCA, Methods), covering the entire chromosome, as indicated in Figure 4b. For each indentation curve, we calculated the viscoelasticity index and Young’s Modulus, and plotted their value as a function of the distance between indentation point and CCA.

**Figure 4:**
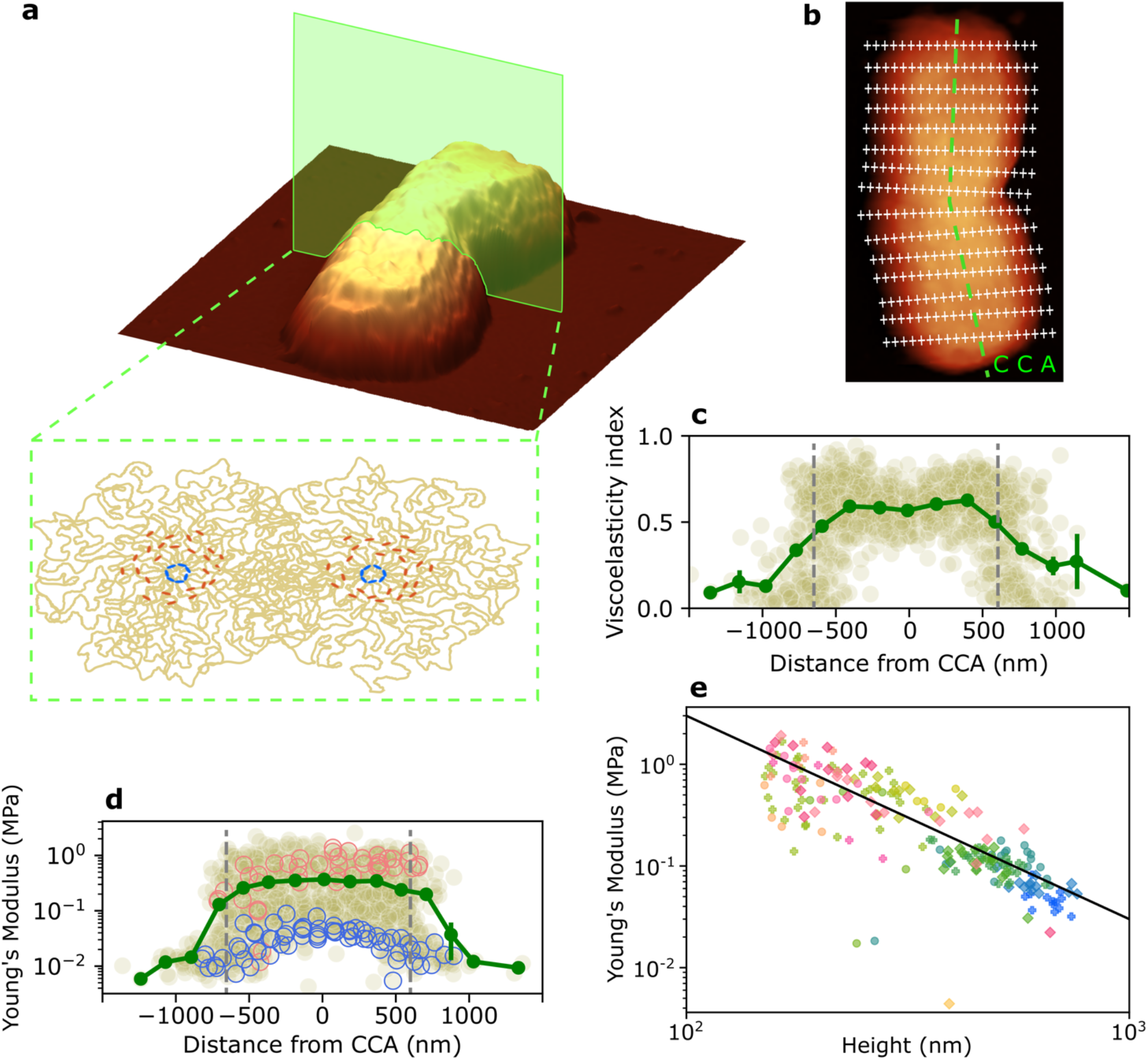
a) Schematic illustration of the chromatids cross section; outer parts of the chromatids display a lower density of chromatin compared to the central axes, where most of the condensin I (red dots) and II (blue dots) are located. An increased chromatin density can be also found where the two sister chromatids overlap, this region is also characterized by the presence of SCIs (displayed in Figure 1c). b) Representative grid describing all the indentations (white crosses) performed on a chromosome; the middle points of each line (calculated as the middle point between the two outermost valid force curves) define the so-called Chromosome Central Axis (CCA), here represented by the green dashed line. The viscoelasticity index and Young’s Modulus were calculated for each point of the grid (for all the chromosomes) and were plotted as a function of the point distance from the CCA in c) and d). The dark green points in c) and d) represent the average values of viscoelasticity index and Young’s Modulus obtained by binning the scatterplots along the x-axis; error bars report the SEM. The gray dashed vertical lines are located at ± 700 nm from the CCA and separate the outermost parts of the chromatids which behave mechanically different from the inner ones. Blue and pink circles in d) identify the Young’s Modulus values of two representative chromosomes. e) Young’s Modulus scales with the height of the chromosomes according to the expression E ∼ h^−2^; markers with different colors and shapes identify values from indentations performed on different chromosomes.

Figure 4c reports the values of viscoelasticity index recorded at different distances from the CCA (for 17 chromosomes). The dark green trace describes the average values of *η* when binned along the x-axis. The lower values of *η* in the outer parts of the chromatids suggest that chromatin-sparser areas respond more elastically to deformations. In contrast, the center of each chromatid is characterized by higher *η* values. This might be due to the higher chromatin and protein density of these areas, which generate a more entangled network that eventually takes longer to recover the initial structure. Finally, in the vicinity of the CCA, where the sister chromatids overlap, we recorded slightly lower *η* values, probably due to the presence of SCIs that increase the elasticity of the chromatin network. Taken together these results highlight that chromatin-dense regions take longer to relax from deformations compared to chromatin-sparse regions. Repeating the same characterization in selected regions along the chromatids (Figure S5 and Methods) yielded very similar results (Figure S6, S7 a) for both centromeric and mid-arm regions (with only minor variations for the peripheral regions).

To investigate whether chromatin density also mediates chromosome stiffness, we calculated the local Young’s Modulus at different distances from the CCA. The scatterplot in Figure 4d shows that regions located within 700 nm from the CCA (delimited by the dashed vertical gray lines and likely containing the central parts of the chromatids) are dramatically stiffer than the external regions, with local Young’s Moduli almost two orders of magnitude higher. The average Young’s Modulus recorded within 700 nm from the CCA is 290 ± 10 kPa (mean ± SEM), comparable with previous measurements on chromosome spreads rehydrated in PBS ^18^. This result could reflect the higher density of chromatin and SCIs of these central regions. Selectively probing centromeric, mid-arm and peripheral regions yielded similar Young’s Moduli with the same two-orders of magnitude variation (Figure S6 and S7 b).

The colored markers (blue and pink) in Figure 4d identify the Young’s Modulus values for two representative chromosomes and highlight how the observed two-orders of magnitude variability can be mostly explained by inter-chromosomal differences. In previous work, we found that the dissipation due to entangled chromatin loops scaled strongly with the chromosome density ρ ^11^. Therefore, we wondered if these differences in Young’s Modulus could also be density dependent. As shown in Figure 4e, we find that the Young’s Modulus scales with *h* (i.e., the contact point) as *E* ∼ *h*^−2^, and that differences in Young’s Modulus between chromosomes can mostly be attributed to their differences in height. Since differences in chromosome widths are relatively small and uncorrelated with differences in height (Figure S8), we argue that *h* is mostly determined by how much each sample is pressed against the substrate and directly reflects the density: ρ ∼ *h*^−1^. Interestingly, our micromechanical results are consistent with the scaling *E* ∼ ρ^2^ for an entangled solution of flexible polymers ^36^, providing further evidence that we are probing the response of entangled chromatin loops surrounding the central scaffold of the chromosome.

## Conclusions

In this study we have shown how AFM can contribute to characterize both the interface structure and the internal mechanics of native human mitotic chromosomes. Different from previous AFM studies^17–19^, we probed chromosomes within a liquid solution mimicking the mitotic environment. Imaging in these conditions allowed capturing features of the chromatin organization over the entire range of length that characterizes chromosome structure. Compared to results from previous structural models^5^, our images suggest that not all the chromatin fibers are arranged in loops. We also probed chromosome surface and found that this interface exhibits self-similarity over a broad range of length scales. This analysis provides a novel perspective for better understanding the properties of the chromosome surface and how intracellular interactions are mediated by this interface during mitosis. With respect to previous OT^15^ and micropipette studies^13,14,37^ that probed the mechanical response of the entire chromosome body, we show that AFM measurements could be used to isolate the sole mechanical contribution of the chromatin network. Comparing results from our microrheology analysis with OT experiments^15^ showed that the chromatin network possesses a fluid-like behavior with energy dissipation one order of magnitude higher than that of the entire chromosome body. Moreover, we found that the relaxation of the chromatin network follows two characteristic timescales which are faster than the ones predicted by previous micropipette experiments^13^ for the stiffer protein-rich scaffold. Finally, AFM-FS revealed that the local mechanics of the chromatin network is mediated by its density and is highly heterogeneous across different chromosomes. Taken together, our results improve the current understanding of mitotic chromosomes by revealing their scale-dependent structure and local mechanics. Moreover, we developed a toolbox of analyses that can be applied to different follow-up investigations on the role of individual molecular components (SMC proteins and surface proteins, such as Ki67) to the observed structure, surface, and mechanics of chromosomes, in both healthy and pathological conditions.

## Methods

All the experiments were carried out in polyamine buffer (PA buffer) containing 15 mM TRIS (buffered at pH = 8), 2 mM EDTA, 80 mM KCl, 20 mM NaCl, 0.5 mM EGTA, 0.2 % Tween20, 0.5 mM spermidine and 0.2 mM spermine. All buffers were made with ultrapure water (MilliQ, Millipore) and filtered (0.2 µm pore size, Whatman) before use.

### Chromosome isolation

Cell culture and chromosome isolation were performed according to procedures already used and described by previous works^15,38^. Briefly, U2OS TRF1-BirA H2B-eGFP cells were cultured in high glucose GlutaMAX DMEM (Gibco), containing 10% FBS (Gibco) and penicillin-streptomycin (Gibco) at 37°C, 5% CO2. 48h prior to chromosome isolation, the medium was supplemented with 12.2 µg/mL biotin. Metaphase arrest was achieved by incubating the cells with 50 ng/mL nocodazole for 4h. Arrested cells were harvested via mitotic shakeoff, pelleted by centrifugation and washed twice with PBS. The mitotic cells were then incubated for 10 min at room temperature in a buffer consisting of 75 mM KCl and 5 mM Tris-HCl pH 8.0. Cells were then resuspended in polyamine buffer (see above) supplemented with Complete mini protease inhibitor and PhosSTOP phosphatase inhibitor (Roche). Using a dounce homogenizer with a tight-fitting pestle, the cells were lysed in ∼25 strokes. The lysate was cleared by centrifugation and added to a glycerol step gradient (60% glycerol followed by 30% glycerol, both in polyamine buffer) and centrifuged at 1750 RCF for 30 minutes. The 60% glycerol fraction, containing the mitotic chromosomes, was stored at -20°C.

### Surface and sample preparation for AFM experiments

All the AFM experiments were performed on poly-L-lysine (PLL)-coated glass coverslips. The coverslips were cleaned in a sonicator bath for 1 hour in 2 % (v/v) Hellmanex solution, followed by 30 minutes in milliQ water; after that, the coverslips were dried with an argon flow and stored in a sealed box. Before each experiment, the glass coverslips were activated with air plasma (Harrick Plasma Expanded Plasma Cleaner 115V) for 3 minutes followed by immediate immersion in milliQ water for other 5 minutes. The activated slides were then incubated for 30 minutes in a 0.01 % (w/v) PLL solution in 0,5 mM, pH 8.15 borate buffer (Na_3_BO_3_), at room temperature; at the end of the incubation the slides were thoroughly rinsed with milliQ water, dried with a gentle argon flow and placed in the AFM for the measurements. A 10μl droplet of the chromosome preparation, diluted 1:10 in the PA buffer, was spotted on the glass coverslip and let to equilibrate for 20 minutes at room temperature before starting the experiments. All the AFM experiments (except for microrheology, see below) were performed on a Bruker Bioscope Catalyst coupled with an inverted optical microscope, at room temperature and using the PA buffer as imaging solution. The AFM was equipped with different probes depending on whether it was used for imaging or Force Spectroscopy (FS) experiments. Each chromosome was first located using the Nikon ECLIPSE Ti-U epi-fluorescence microscope, equipped with a 10x objective. After that, the AFM was used to scan the region of interest (ROI) and probe the chromosome with higher spatial accuracy and, in the case of AFM-FS, to run the mechanical characterization.

### AFM imaging

Images were collected in PeakForce Tapping mode, using an Olympus BLAC40TS probe; the force set point and the other imaging parameters were tuned in order to accurately follow the topological features of the chromosome structure without introducing any deformation. Images were processed using Gwyddion 2.58^39^.

#### Chromosome surface roughness analysis

The height profiles displayed in Figure S2 were obtained using Gwyddion 2.58 and saved as .txt file for further analyses. For each profile, the average standard deviation is calculated over a rolling sampling window. The size of the sampling window converted in real units represents the length scale (*L*) under consideration. By repeating this procedure for different window sizes and by averaging the standard deviations obtained from all the profiles, we obtain the plot in Figure 1f. A power-law fit (weighted on the variability of each value in the plot) is applied to the portion of the plots presenting 40 nm < *L* < 250 nm to yield the so-called scaling exponent. The entire procedure was executed using a custom Python script.

#### Chromatin halo height distribution

Gwyddion 2.58^39^ was used to extract the height data of the chromatin halo (masked area in Figure S1). The height distribution was fitted with a sum of 5 Gaussian curves using a custom Matlab script.

### AFM-Force Spectroscopy (AFM-FS)

DNPS10 probes (cantilever C, nominal tip radius 10 nm, nominal elastic constant 0.24 N/m, from Bruker, Camarillo, CA, USA), calibrated with the thermal noise method^40^ were used for all the AFM-FS measurements. Images at lower resolution were used to localize each chromosome in the center of the ROI; after that, multiple indentations (ramp size 1 μm and indentation rate 0.5 Hz) were performed on different parts of the chromosome. At the end of the indentations, a second image was performed to ensure that no permanent alterations were induced to the chromosome morphology. We note however, that this happened very rarely. The recorded force curves were processed with custom made Python scripts to extract the information about the mechanical response of mitotic chromosomes. For each curve, the contact point was determined as the first data point along the indenting force curve (green trace in Figure 2) at which a force greater than 50 pN was recorded (this threshold was chosen based on the signal to noise ratio that characterized the collected force curves). To ensure considering only indentations performed on the chromosome body, force curves presenting a contact point < 150 nm were discarded from the analysis. For extracting the value of Young’s Modulus from each force curve, the modified Hertz fit from Dimitriadis et al.^28^ was applied to the indenting curve; more precisely, each curve was fitted from the contact point to an indentation depth equal to 30% of the chromosome thickness (estimated from the value of the contact point). The viscoelasticity index, *η*, is related to the energy dissipated during the indentation process and is defined as *η* = 1 − *A*_2_⁄*A*_1_, where *A*_1_and *A*_2_ represent the areas under the indenting and retracting curves, calculated within the loading and unloading regimes (i.e., where the tip-sample interactions result in positive force values), respectively^29,30^. A custom Python script was used to calculate the areas under the indenting and retracting curves and to compute the values of *η*.

#### Cross-chromatid analysis

Series of indentations along lines distributed perpendicularly to the chromosome long axis and covering the entire chromosome body, as described in Figure 4b were performed on a total of 17 chromosomes. Indentations showing a contact point < 150 nm were declared 1non valid1 and discarded from the analysis to ensure considering only indentations probing the actual chromosome body. For each indentation line, the coordinates of the middle point between the two most distal valid indentations were calculated; the chromosome central axis (CCA) was defined by all the middle points obtained via this procedure. The results shown in Figure 4c, d report the results for all the valid indentations on all the probed chromosomes. The analysis was carried out using custom Python scripts.

#### Cross-chromatid analysis of centromeric, mid-arm and peripheral regions

We selected three different regions along the chromatids: the centromeric regions (referred as centromeric), the middle part of each arm (referred as mid-arm) and the most distal part of the arms (referred as peripheral), as shown in Figure S5. Despite the length of the telomeres in chromosomes from U2OS cells can reach values > 50 kb^41^, these values are still relatively small compared to the areas probed in the peripheral regions by the AFM tip; for this reason, we consider the indentations performed on the peripheral regions more representative of the subtelomeric structures^42^. For each region, we performed series of indentations along three lines running perpendicularly to the chromosome long axis (Figure S5); a total of 21 chromosomes were probed. *Mid-arm* regions were selected at approximately half-distance between the centromere and each chromatid end. Due to diversity in chromosome arm’s length, it was possible to probe the arms on both sides of the centromere for only 6 chromosomes; for the other 15, only one side of the centromere presented arms that were long enough to enable a clear distinction between peripheral and centromeric regions. For all the 21 chromosomes it was possible to probe the peripheral regions (on both sides of the centromere) and the centromeric region. For each chromosome, the three different regions were located arbitrarily, since the instrumentation did not allow for a rigorous distance parametrization among the different chromosomes. Analysis of the indentation curves was performed using the same procedure described in the Cross-chromatids analysis, using custom Python scripts.

#### Comparing chromosome widths to heights

To determine whether there is any correlation between the width and the height of sampled chromosomes (the 17 chromosomes from the cross-chromatids analysis), we compared their mean widths and heights (Figure S8). To obtain values for the mean width, three height profiles were traced for each chromosome in parts of the chromatids that exhibited a uniform width. For each profile, the width of the chromatids was defined as the width of the region higher than 150 nm, to discard the chromatin halo surrounding the chromosomes. The mean height values for each chromosome were obtained by averaging the contact point heights for all indentations performed within 200 nm from the CCA.

#### Young’s Modulus scaling with chromosome height

For determining whether the compaction of a chromosome affects its Young’s modulus, we used data from all measurement points within 200 nm from the CCA. For each measurement point, we plotted the Young’s modulus against the contact point height (Figure 4e).

### Microrheology measurements

Microrheology experiments were performed using Dynamic mechanical analysis (DMA) on a Bruker NanoWizardV BioAFM. Surface and sample preparations were identical to the ones used for the other AFM-FS experiments. A modified version of the PFQNM-LC-A-CAL probe, carrying a cylindrical tip with end radius of 870 nm and spring constant 0.137 N/m (Bruker, Camarillo, CA, USA) was used for the measurements. The commercially available version is now released as SAA-SPH-1UM (Bruker, Camarillo, CA, USA). The AFM setup was coupled with a fluorescence microscope that was used to locate the eGFP-labelled chromosomes on the surface. Once a single chromosome was localized within the ROI, multiple force curves were performed over the chromosome body. For these experiments, each measurement consisted of four different steps (see Figure 3a): i) a shallow penetration of the sample until reaching a force of 150 pN, ii) a relaxation phase in which the tip was kept at a constant height and the relaxation force was monitored, iii) an oscillatory part where oscillatory deformations (with 10 nm amplitude) were applied to the sample and iv) a final retraction phase in which the tip was brought back to its initial position. For each chromosome, three different oscillatory frequencies were probed: 2, 20 and 200 Hz. Data were collected on 19 chromosomes. At least 15 force curves were collected from each chromosome for each oscillatory frequency. The FRC analysis was performed only on the dataset related to the 20 Hz oscillation frequency; this arbitrary decision is motivated by the fact that the parameters used in the 1Approach1 and 1Relaxation1 parts of the force curves (Figure 3a) did not change when probing different oscillation frequencies. The data were analyzed combining the proprietary *JPK-SPM Software* (version 8.0.74) with custom Python scripts (for plotting and fitting purposes).

## Acknowledgements

We acknowledge Dr. Alexander Dulebo and Bruker (Berlin, Germany) for supporting the microrheology studies and for the use of the Bruker NanoWizardV BioAFM. We acknowledge Ian Hickson (Center for Chromosome, Department of Cellular and Molecular Medicine, University of Copenhagen, Denmark) for fruitful discussions and help in developing the chromosome isolation procedure. This research was supported by the European Research Council under the European Union’s Horizon 2020 research and innovation program (MONOCHROME, grant agreement no. 883240 to G.J.L.W., and Learn4DChromosome, Grant agreement No. 101122863 to C.P.B.).

## References

1. Paulson, J. R., Hudson, D. F., Cisneros-Soberanis, F. & Earnshaw, W.C. Mitotic chromosomes. Semin. Cell Dev. Biol. 117, 7–29 (2021).

2. Spicer, M. F. D. & Gerlich, D. W. The material properties of mitotic chromosomes. Curr. Opin. Struct. Biol. 81, 102617 (2023).

3. Schneider, M. W. G. et al. A mitotic chromatin phase transition prevents perforation by microtubules. Nature 609, 183–190 (2022).

4. Man, T., Witt, H., Peterman, E. J. G. & Wuite, G. J. L. The mechanics of mitotic chromosomes. Q. Rev. Biophys. 54, e10 (2021).

5. Gibcus, J. H. et al. A pathway for mitotic chromosome formation. Science 359, (2018).

6. Nishino, Y. et al. Human mitotic chromosomes consist predominantly of irregularly folded nucleosome fibres without a 30-nm chromatin structure. EMBO J. 31, 1644–1653 (2012).

7. Chicano, A. et al. Frozen-hydrated chromatin from metaphase chromosomes has an interdigitated multilayer structure. EMBO J. 38, 1–12 (2019).

8. Beel, A. J., Azubel, M., Matteï, P.-J. & Kornberg, R. D. Structure of mitotic chromosomes. Mol. Cell 81, 4369-4376.e3 (2021).

9. Walther, N. et al. A quantitative map of human Condensins provides new insights into mitotic chromosome architecture. J. Cell Biol. 217, 2309–2328 (2018).

10. Kubalová, I. et al. Helical coiling of metaphase chromatids. Nucleic Acids Res. 51, 2641–2654 (2023).

11. Witt, H. et al. Ion-mediated condensation controls the mechanics of mitotic chromosomes. 2023.04.11.536423 Preprint at 10.1101/2023.04.11.536423 (2023).

12. Beel, A. J., Matteï, P.-J. & Kornberg, R. D. Mitotic Chromosome Condensation Driven by a Volume Phase Transition. http://biorxiv.org/lookup/doi/10.1101/2021.07.30.454418 (2021) doi:10.1101/2021.07.30.454418.

13. Poirier, M. G., Nemani, A., Gupta, P., Eroglu, S. & Marko, J. F. Probing Chromosome Structure with Dynamic Force Relaxation. Phys. Rev. Lett. 86, 360–363 (2001).

14. Sun, M., Biggs, R., Hornick, J. & Marko, J. F. Condensin controls mitotic chromosome stiffness and stability without forming a structurally contiguous scaffold. Chromosome Res. 26, 277–295 (2018).

15. Meijering, A. E. C. et al. Nonlinear mechanics of human mitotic chromosomes. Nature 605, 545– 550 (2022).

16. Krieg, M. et al. Atomic force microscopy-based mechanobiology. Nat. Rev. Phys. (2018) doi:10.1038/s42254-018-0001-7.

17. Nomura, K. et al. Visualization of elasticity distribution of single human chromosomes by scanning probe microscopy. Jpn. J. Appl. Phys. Part 1 Regul. Pap. Short Notes Rev. Pap. 44, 5421– 5424 (2005).

18. Jiao, Y. & Schäffer, T. E. Accurate Height and Volume Measurements on Soft Samples with the Atomic Force Microscope. Langmuir 20, 10038–10045 (2004).

19. Ushiki, T. & Hoshi, O. Atomic force microscopy for imaging human metaphase chromosomes. Chromosome Res. 16, 383–396 (2008).

20. Kalle, W. & Strappe, P. Atomic force microscopy on chromosomes, chromatin and DNA: A review. Micron 43, 1224–1231 (2012).

21. Lipiec, E. et al. Infrared nanospectroscopic mapping of a single metaphase chromosome. Nucleic Acids Res. 47, e108 (2019).

22. Roh, S. et al. Direct observation of surface charge and stiffness of human metaphase chromosomes. Nanoscale Adv. 5, 368–377 (2023).

23. Pfosser, M., Königshofer, H. & Kandeler, R. Free, Conjugated, and Bound Polyamines during the Cell Cycle of Synchronized Cell Suspension Cultures of Nicotiana tabacum. J. Plant Physiol. 136, 574–579 (1990).

24. Sen, N. et al. Physical Proximity of Sister Chromatids Promotes Top2-Dependent Intertwining. Mol. Cell 64, 134–147 (2016).

25. Eltsov, M., MacLellan, K. M., Maeshima, K., Frangakis, A. S. & Dubochet, J. Analysis of cryo-electron microscopy images does not support the existence of 30-nm chromatin fibers in mitotic chromosomes in situ. Proc. Natl. Acad. Sci. 105, 19732–19737 (2008).

26. Mandelbrot, B. B., Passoja, D. E. & Paullay, A. J. Fractal character of fracture surfaces of metals. Nature 308, 721–722 (1984).

27. Pande, C. S., Richards, L. E., Louat, N., Dempsey, B. D. & Schwoeble, A. J. Fractal characterization of fractured surfaces. Acta Metall. 35, 1633–1637 (1987).

28. Dimitriadis, E. K., Horkay, F., Maresca, J., Kachar, B. & Chadwick, R. S. Determination of elastic moduli of thin layers of soft material using the atomic force microscope. Biophys. J. 82, 2798– 2810 (2002).

29. Briscoe, B. J., Fiori, L. & Pelillo, E. Nano-indentation of polymeric surfaces. J. Phys. Appl. Phys. 31, 2395–2405 (1998).

30. Klymenko, O., Wiltowska-Zuber, J., Lekka, M. & Kwiatek, W. M. Energy Dissipation in the AFM Elasticity Measurements. Acta Phys. Pol. A 115, 548–551 (2009).

31. Alcaraz, J. et al. Microrheology of Human Lung Epithelial Cells Measured by Atomic Force Microscopy. Biophys. J. 84, 2071–2079 (2003).

32. Doi, M. & Edwards, S. F. The Theory of Polymer Dynamics. (Clarendon Press, Oxford, 1986).

33. Dey, A., Shi, G., Takaki, R. & Thirumalai, D. Structural changes in chromosomes driven by multiple condensin motors during mitosis. Cell Rep. 42, 112348 (2023).

34. Gironella-Torrent, M., Bergamaschi, G., Sorkin, R., Wuite, G. & Ritort, F. Viscoelastic phenotyping of red blood cells. Biophys. J. S0006349524000353 (2024) doi:10.1016/j.bpj.2024.01.019.

35. Marsden, M. P. F. & Laemmli, U. K. Metaphase chromosome structure: Evidence for a radial loop model. Cell 17, 849–858 (1979).

36. Gennes, P.G.de. Remarks on entanglements and rubber elasticity. J. Phys. Lett. 35, 133–134 (1974).

37. Poirier, M., Eroglu, S., Chatenay, D. & Marko, J. F. Reversible and irreversible unfolding of mitotic newt chromosomes by applied force. Mol. Biol. Cell 11, 269–276 (2000).

38. Clement, T. V. M., van der Smagt, C. & Wuite, G. J. L. Probing Mitotic Chromosome Mechanics Using Optical Tweezers. in Single Molecule Analysis : Methods and Protocols (eds. Heller, I., Dulin, D. & Peterman, E.J.G.) 91–107 (Springer US, New York, NY, 2024). doi:10.1007/978-1-0716-3377-9_5.

39. Nečas, D. & Klapetek, P. Gwyddion: An open-source software for SPM data analysis. Cent. Eur. J. Phys. 10, 181–188 (2012).

40. Hutter, J. L. & Bechhoefer, J. Calibration of atomic-force microscope tips. Rev. Sci. Instrum. 64, 1868–1873 (1993).

41. Ma, H., Reyes-Gutierrez, P. & Pederson, T. Visualization of repetitive DNA sequences in human chromosomes with transcription activator-like effectors. Proc. Natl. Acad. Sci. 110, 21048– 21053 (2013).

42. Mefford, H. C. & Trask, B. J. The complex structure and dynamic evolution of human subtelomeres. Nat. Rev. Genet. 3, 91–102 (2002).

