## Supplementary Figures and Data for "Resolving interface structure and local internal mechanics of mitotic chromosomes"

### Supplementary Information

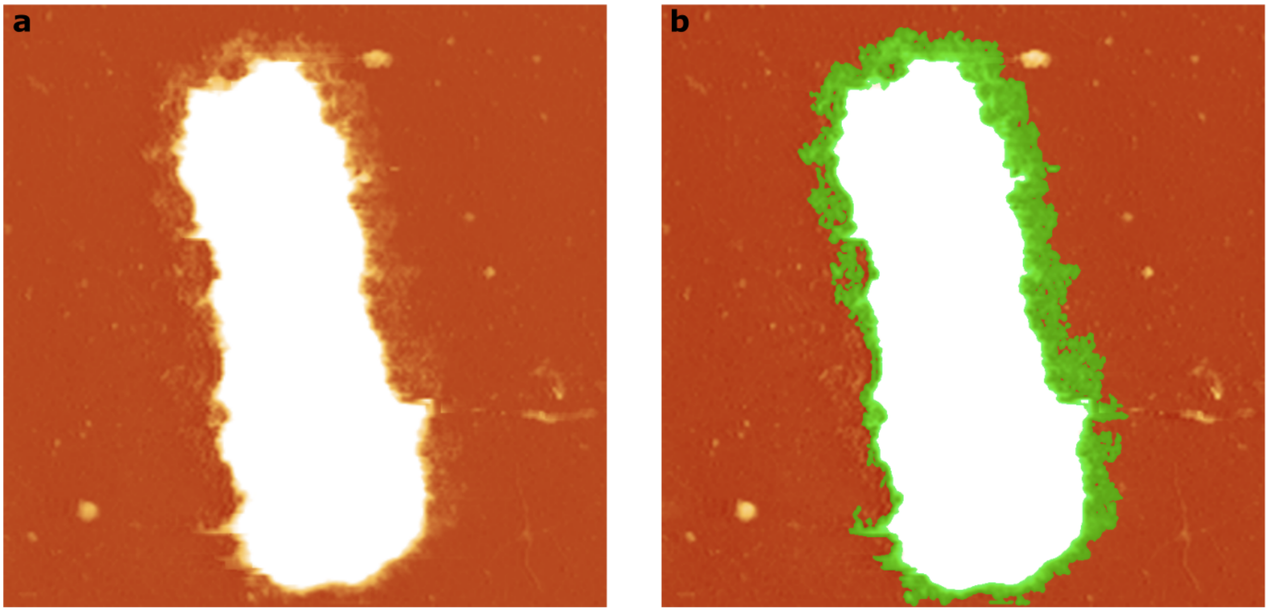


***Figure S1:*** *a) Increasing the contrast of Figure 1a reveals the presence of chromatin fibers and loops protruding outwards from the entire chromosome body. b) The green area represents the selected regions that were used to obtain the height distribution in Figure 1e.*

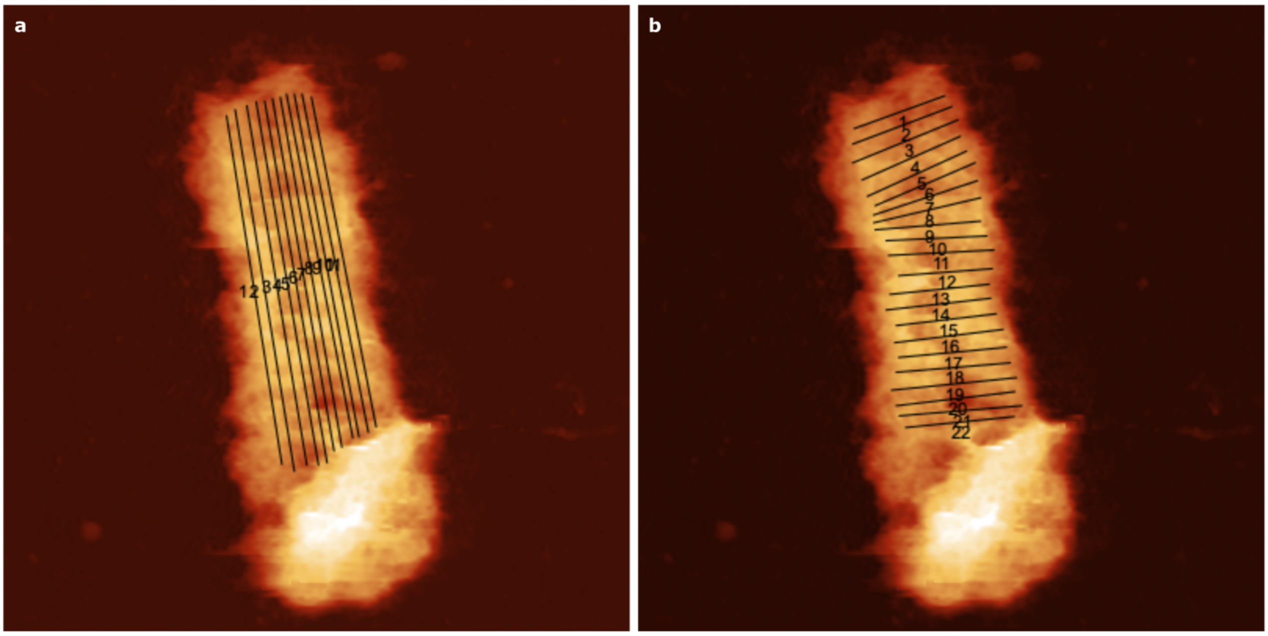


***Figure S2:*** *a) Longitudinal height profiles used to obtain the plot in Figure 1g. b) Height profiles running perpendicular to the chromosome long axis, used to obtain the plot in Figure S3.*

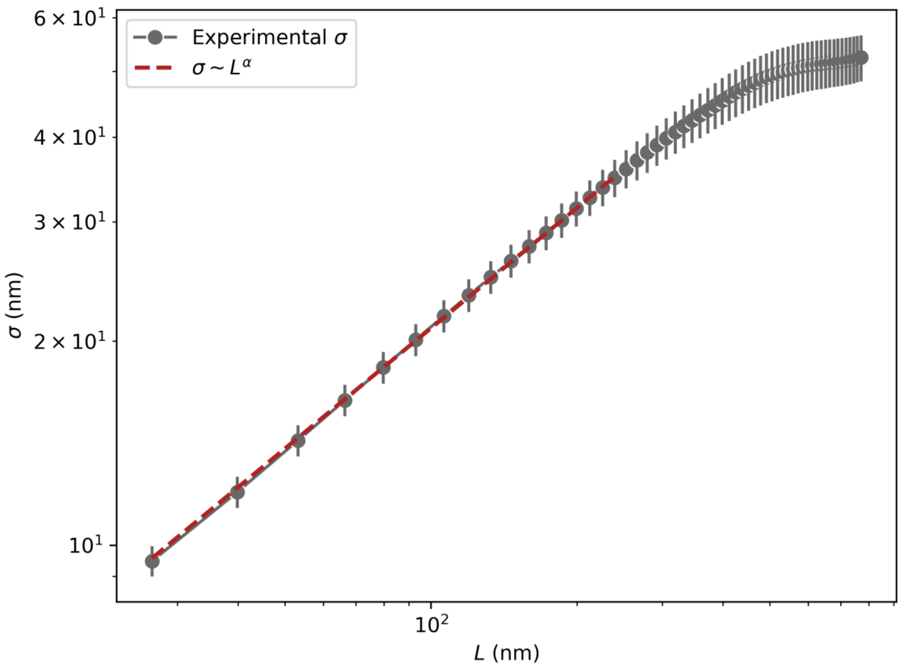


***Figure S3:*** *Average roughness as a function of different length scales L for height profiles running perpendicularly to the chromosome long axis (Figure S2 b), refer to main text and Methods for details about fitting and scaling exponent.*

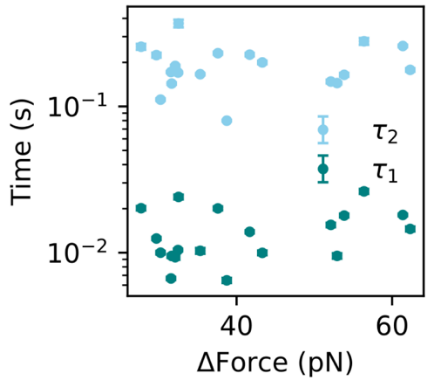


***Figure S4:*** *Values of the two characteristic timescales obtained from the fits of the FRCs in Figure 4g, plotted as a function of the respective force drop, calculated as the difference between the forces at the start of the relaxation and the values of* $F_r$ *predicted by the fits (error bars report the SEM).*

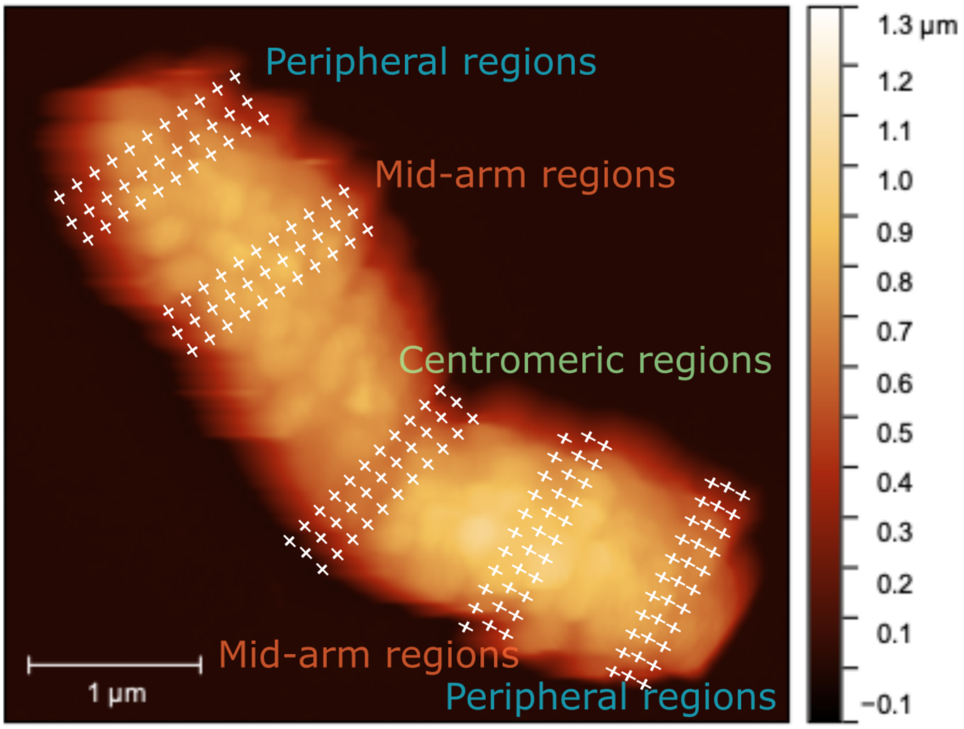


***Figure S5:*** *Schematics showing the experimental design for the indentations performed on different regions along the chromatids. For each region (peripheral, centromeric and mid-arm), we performed indentations along 3 different lines running perpendicularly to the CCA. If possible, peripheral and mid-arm regions were probed for both arms. Results obtained from all the chromosomes were grouped together based on the indented region.*

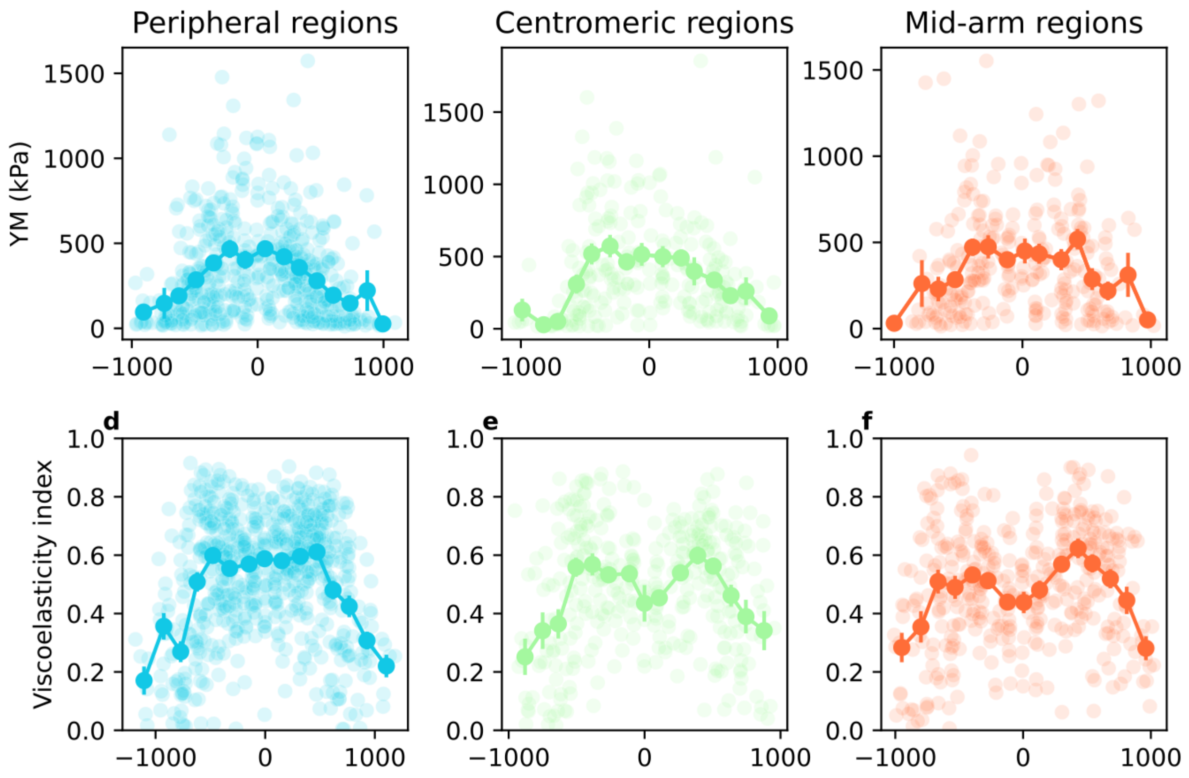


***Figure S6:*** *Scatterplots of the Young's Modulus and viscoelasticity index obtained from indentations on different regions along the chromatids. Darker traces report the average values binned along the x-axis (N=20 chromosomes, error bars report the SEM)*

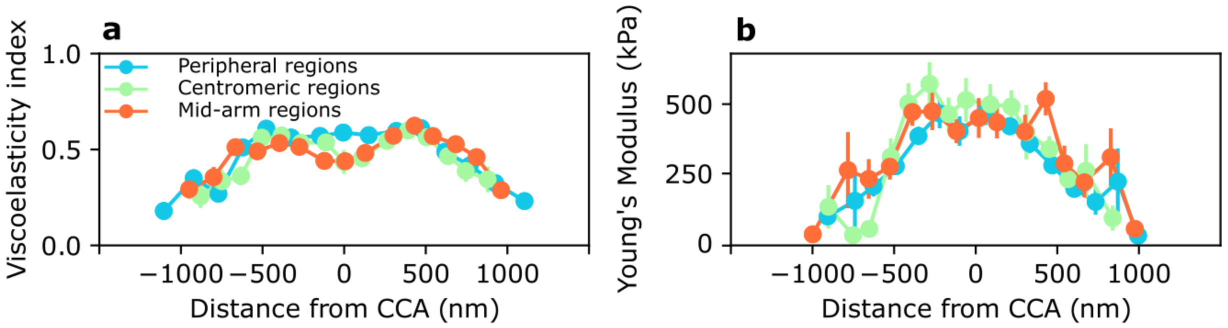


***Figure S7:*** *Average values binned along the x-axis of Figure S6 reported together for a better comparison. a) viscoelasticity index profiles for centromeric and mid-arm regions are similar, peripheral regions behave slightly different in the vicinity of CCA, possibly due to the presence of less SCIs. b) Young's Modulus profiles for peripheral, centromeric and mid-arm regions show the same trend, varying over two orders of magnitude over the entire width of the chromatids.*

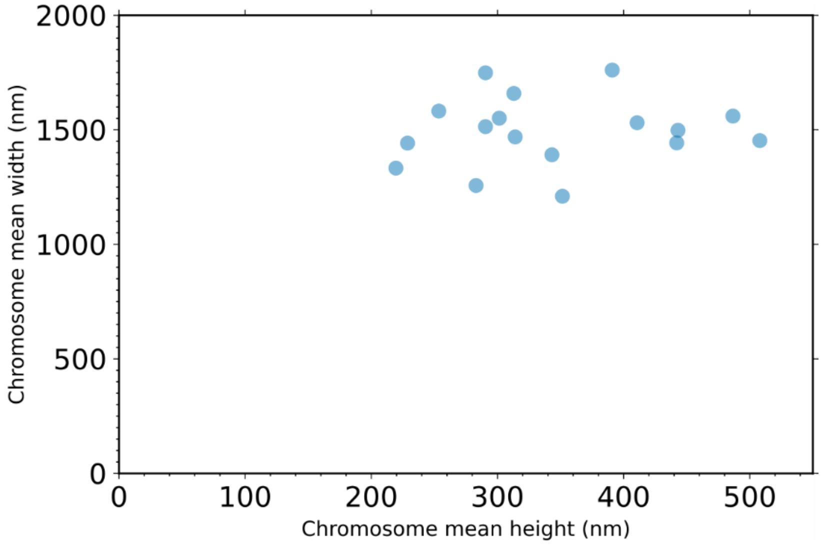


***Figure S8:*** *Mean values of height and width for the 17 chromosomes analyzed in Figure 4, showing weak/absent correlation between the two properties (refer to Methods section for more details).*

***Table S1: Results of the fits to the FRCs***

| *Chrm #* | $\tau_1$ *(s)* | $\tau_1$ *std* | $\tau_2$ *(s)* | $\tau_2$ *std* | $F_r$ *(pN)* | $F_0$ *(pN)* | *Force drop (pN)* |
|---|---|---|---|---|---|---|---|
| *1* | 0.0201 | 0.0006 | 0.2311 | 0.0050 | 114.3784 | 151.9811 | 37.6026 |
| *2* | 0.0139 | 0.0003 | 0.2260 | 0.0057 | 111.9847 | 153.6741 | 41.6894 |
| *3* | 0.0201 | 0.0007 | 0.2559 | 0.0117 | 88.8980 | 116.6186 | 27.7206 |
| *4* | 0.0240 | 0.0009 | 0.3694 | 0.0221 | 85.2793 | 117.8226 | 32.5433 |
| *5* | 0.0181 | 0.0003 | 0.2589 | 0.0046 | 91.1969 | 152.5780 | 61.3811 |
| *6* | 0.0263 | 0.0010 | 0.2787 | 0.0145 | 97.1481 | 153.5396 | 56.3915 |
| *7* | 0.0103 | 0.0005 | 0.1660 | 0.0025 | 102.6126 | 137.9421 | 35.3295 |
| *8* | 0.0065 | 0.0003 | 0.0799 | 0.0010 | 111.0176 | 149.7605 | 38.7429 |
| *9* | 0.0095 | 0.0004 | 0.1444 | 0.0029 | 95.4760 | 148.4048 | 52.9288 |
| *10* | 0.0179 | 0.0005 | 0.1645 | 0.0047 | 93.0978 | 146.9372 | 53.8394 |
| *11* | 0.0155 | 0.0005 | 0.1479 | 0.0042 | 99.4077 | 151.5301 | 52.1224 |
| *12* | 0.0145 | 0.0006 | 0.1778 | 0.0047 | 78.8625 | 141.1926 | 62.3301 |

| | | | | | | | |
|---|---|---|---|---|---|---|---|
| *13* | 0.0100 | 0.0004 | 0.1998 | 0.0053 | 107.3481 | 150.6597 | 43.3116 |
| *14* | 0.0095 | 0.0003 | 0.1433 | 0.0023 | 123.8626 | 155.5212 | 31.6586 |
| *15* | 0.0125 | 0.0003 | 0.2243 | 0.0084 | 124.0319 | 153.7253 | 29.6934 |
| *16* | 0.0100 | 0.0003 | 0.1113 | 0.0021 | 118.9525 | 149.1795 | 30.2271 |
| *17* | 0.0067 | 0.0002 | 0.1718 | 0.0024 | 115.6326 | 147.2018 | 31.5692 |
| *18* | 0.0093 | 0.0004 | 0.1891 | 0.0034 | 115.1471 | 147.2601 | 32.1130 |
| *19* | 0.0104 | 0.0002 | 0.1703 | 0.0023 | 108.8279 | 141.3020 | 32.4740 |
